# Cotranslational assembly imposes evolutionary constraints on homomeric proteins

**DOI:** 10.1101/074963

**Authors:** Eviatar Natan, Tamaki Endoh, Liora Haim-Vilmovsky, Guilhem Chalancon, Tilman Flock, Jonathan TS. Hopper, Bálint Kintses, Lejla Daruka, Gergely Fekete, Csaba Pál, Balázs Papp, Peter Horvath, Joseph A. Marsh, Adrian H. Elcock, M Madan Babu, Carol V. Robinson, Naoki Sugimoto, Sarah A. Teichmann

## Abstract

There is increasing evidence that some proteins fold during translation, *i.e.* cotranslationally, which implies that partial protein function, including interactions with other molecules, could potentially be unleashed early on during translation. Although little is known about cotranslational assembly mechanisms, for homomeric protein complexes, translation by the ribosome, folding and assembly, should be well-coordinated to avoid misassembly in the context of polysomes. We analysed 3D structures of homomers and identified a statistically significant trend conserved across evolution that supports this hypothesis: namely that homomeric contacts tend to be localized towards the C-terminus rather than N-terminus of homomeric polypeptide chains. To probe this in more detail, we expressed a GFP-based library of 611 homomeric *E. coli* genes, and analyzing their folding and assembly *in vivo*. Consistent with our hypothesis, interface residues tend to be located near the N-terminus in cotranslationally aggregating homomers. In order to dissect the mechanisms of folding and assembly under controlled conditions, we engineered a protein library with three variable components: (i) the position and type homomerization domain, (ii) the reporter domain and (iii) the linker length that connects the two. By analyzing the misassembly rates of these engineered constructs *in vivo*, *in vitro* and *in silico*, we confirmed our hypothesis that C-terminal homomerization is favorable to N-terminal homomerization. More generally, these results provide a set of spatiotemporal constraints within polypeptide chains that favor efficient assembly, with implications for protein evolution and design.

## Introduction

Early in protein synthesis, a nascent chain begins to sprout from the ribosome’s exit tunnel into the crowded cellular milieu. There is increasing evidence that the nascent chain can fold concomitantly with translation, in a process known as cotranslational folding (Elcock, 2006; Pechmann and Frydman, 2013; Sander et al., 2014). Cotranslational folding is thought to have evolved to protect nascent chains from non-specific interactions with the cell content, or from entanglement with other nascent chains in the highly crowded environment of the polysome, which is the super-complex comprised of multiple ribosomes on a single mRNA (Hartl and Hayer-Hartl, 2009).

While cotranslational folding can protect proteins from aggregation, it may also harbor a risk for homomers. Homomers are protein complexes comprised of multiple identical subunits. They are extremely common in the proteomes of all organisms and are involved in all major cellular functions due to the advantages of symmetry, stability and allosteric properties of their quaternary structure (Goodsell and Olson, 2000; Levy and Teichmann, 2013). In particular, the propensity for homomeric self-assembly is significantly enriched in bacterial proteins (Marsh et al., 2015). If a homomeric subunit folds cotranslationally to expose its interface to other nascent chains, the complex may also assemble cotranslationally (Shieh et al., 2015; Wells et al., 2015). This would automatically lead to an increase in the already high local concentration of the nascent chains attached to the same polysome. Therefore, the cotranslational assembly of domains that mediate homomer formation forces unfolded, partially folded or freshly folded parts of the nascent chains into extremely close proximity as they emerge from the ribosome (Figure 1C), which can increase the likelihood of misassembly.

**Figure 1.**
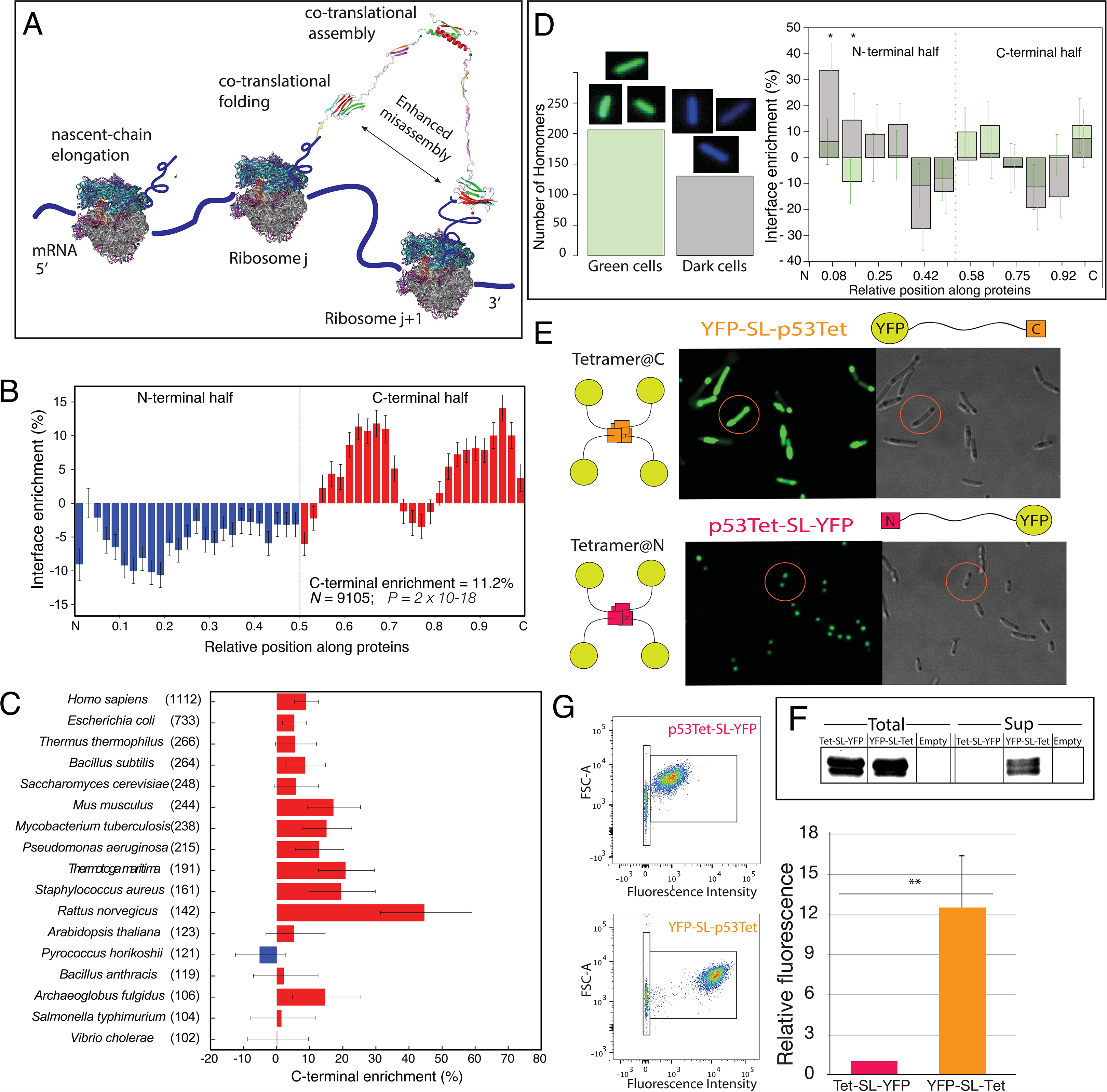
Homomeric interface residues are C-terminally enriched, supporting cotranslational assembly of homomers. (A-B) Distribution of interface-forming residues in the N- vs C-terminal halves of homomeric proteins. (A) Relative enrichment of interface-forming residues along the lengths of homomers. Residues are binned according to their position along the full-length protein (N terminus is 0, the C terminus is 1). The interface enrichment is defined as the proportion of all interfaces present in each bin divided by the proportion of monomer accessible surface area present in that bin. Thus the percentage plotted represents the relative probability that a given point on the surface of a monomer will participate in a homomeric interface. The C-terminal enrichment indicates the overall enrichment in the entire C-terminal half *vs*. the N-terminal half. Error bars represent standard error calculated from 10^6^ bootstrapping replicates, and the *P*-value is derived from the fact that there was a net enrichment in C-terminal interface for all replicates. (B) Relative enrichment in interface in the C-terminal halves of proteins compared to the N-terminal halves for all species with >100 non-redundant homomer structures in our dataset. Error bars represent standard error calculated from 10^4^ bootstrapping replicates *per* species. The number of homomer structures *per* species is given in parentheses. (C) Scheme of the hypothesis. Cotranslational assembly of proteins is intra-molecular rather than inter-molecular. Translation of homomeric proteins within a polysome may lead to misassembly upon cotranslational folding and assembly of two or more nascent-chains. (D) Image-based high-throughput screen reveals an N-terminal enrichment of interface residues in cotranslationally aggregating homomers. On the left, the distribution of *in vivo* solubility states of native *E. coli* homomers (see also Figure S3). ‘Green cells’ are cells that present homogeneously high fluorescence level. ‘Dark cells’ are cells that present background fluorescence levels. On the right, the relative enrichment of interface-forming residues along the protein length for ‘Green cells’ (green) and ‘Dark cells’ (grey) is shown. The latter is significantly enriched in N-terminal interface-forming residues compared to the former. *p*-values based on bootstrap hypothesis testing are 0.022 and 0.038 in the first and second bins, respectively. Bars indicate mean ± standard error. Interface enrichment is calculated as in (B), except that standard errors were calculated based on 10^4^ bootstrapping replicates. The overall C-terminal enrichment is 6.67% and −9.83% for ‘Green’ and ‘Dark cells’, respectively. (E-G) *In vivo* expression of the constructs shows that the position of the oligomerization domain is crucial for the solubility of the expressed protein. (E) Confocal microscopy images of p53Tet-SL-YFP and YFP-SL-p53Tet, with YFP reporter gene. The latter presented homogenous fluorescence throughout the cell, while p53Tet-SL-YFP formed inclusion bodies. (F) A Western Blot shows that the expression level of both constructs was similar, yet only YFP-SL-p53Tet was present in the soluble fraction, meaning the origin of the phenotype lies in a post-transcriptional event. The full gel can be found in Figure S4B. (G) Quantification of fluorescence levels of each strain using Flow Cytometry. The black line on top of the dot-plot indicates the intensity of p53Tet-SL-YFP. Fluorescence Intensity of the protein, which correlates with correct folding, is on the x-axis (** *p*-value <0.01).

For example, incompletely folded identical protein domains close to each other could lead to native-type interactions across different chains instead of correct folding of each domain, resulting in a “domain-swap” scenario of entanglement of chains and misassembly. Analogously, chains consisting of repeated domains with high sequence similarity have been shown to misfold through intermingling, domain-swap type residue-residue interactions (Borgia et al., 2011). In these multi-domain proteins, the close proximity of similar domains folding during translation increases the risk of misassembly, which may have an impact on evolution of proteins with domain-repeats (Wright et al., 2005).

Furthermore, the importance of sufficient folding time prior to the exposure of the unfolded nascent chain to the cellular environment has also been well studied in the context of misassembly propensity as a function of translation rate (Nissley and O'Brien, 2014). In this work, the importance of a stably folded domain prior to its exposure to the next translated domain was found to be crucial. We hypothesize that evolutionary constraints must exist to fine tune the balance between cotranslational folding and assembly in homomeric proteins. Specifically, we speculate that as a general role, the position of interface-forming residues will be an important determinant of misassembly, since the N-terminal regions will be translated first and have a greater chance of cotranslational interaction prior to the completion of correct folding.

To address this, we first calculated the relative position of the assembly interface of thousands of homomers of known three-dimensional structure from the Protein DataBank. It was clear from these calculations that the residues that participate in the homomeric interface are more enriched toward the C-*versus* N-terminus. Importantly, this enrichment is conserved across all branches of the tree life suggesting a fundamental biophysical and molecular biological mechanism.

As a next step, we used a novel high-throughput screen to systematically investigate whether the position of the homomeric interface affects protein stability. The screen employed all homomers of *Escherichia coli* of known three-dimensional, encompassing 611 homomers in total. For each homomer, a GFP reporter was attached to its C-terminus (Kitagawa et al., 2005). This type of GFP labeling is a reliable indicator of the aggregation state of the protein (Waldo et al., 1999). We quantified GFP fluorescence within each *E. coli* cell with bespoke image processing software. This quantitative image processing approach allowed us to distinguish early (possibly cotranslational) aggregation, resulting in no GFP fluorescence, *versus* late post-translational aggregation, leading to fluorescent inclusion bodies. The results of the screen indicate that early and severe aggregation correlates with N-terminal enrichment of the homomer interface residues. This supports our hypothesis that folding and assembly can be linked during translation, which exerts selection pressure on homomer interface residue to be biased towards C- rather than N-terminal positions.

To understand in more detail the mechanism underlying this N- *versus* C-terminal bias, we built a library of constructs that reflects the different characteristics of homomers that are at the heart of the hypothesis. Each construct is comprised of three components:

i. An oligomerization-domain, namely the p53 tetramerization domain (p53Tet), a short domain natively located at the C-terminus, which folds and assembles ultrafast (Mateu et al., 1999). It is therefore not surprising that the domain was found to assemble cotranslationally (Nicholls et al., 2002). We placed the p53Tet either at the N- or C-terminus of our constructs. Remarkably, single amino-acid substitutions in this small domain determine whether the domain forms tetramers, dimers, or stays monomeric in its folded state (Gaglia et al., 2013; Lomax et al., 1998). Here, we study all three oligomeric states.
ii. The second component of our constructs is the reporter domain, for which we included YFP, two versions of GFP and luciferase. These reporters are comprised of a few hundreds amino acids (the sequences are provided in Supplementary Methods), which means that the p53Tet has sufficient time of almost a minute to fold and assemble before the reporter has the opportunity to become fully translated, considering that bacterial translation rates are 10-20 aa/sec (Iwasaki and Ingolia, 2016). The reporter domains were chosen based on detectable signal and rate of folding. The library is divided into sub-libraries according to the reporter domain and the linker length (Figure S4A and Table S1).
iii. The third component of the constructs is the linker to separate the oligomerization-domain and the reporter domain in a spatiotemporal manner. The linker was designed to be flexible and of diverse lengths controling p53Tet local-concentration, *e.g.* at the timepoint that the first amino acids of the reporter exist the ribosome tunnel. Moreover, if assembly took place, the distance between the assembly site and newly translated polypeptide chain of the reporter, are proportional to the linker length. Three different linkers were used, short linker (SL), medium linker (ML) and long linker (LL).

We used this library to probe the assembly of the p53 oligomerization-domain variants both *in vivo* and *in vitro*. These experiments provide a comprehensive framework for understanding the constraints on cotranslational assembly of homomers, specifically the balance between folding and assembly, as well as the role of chaperones. The work was complemented by *in silico* simulations, which help to clarify the mechanism for cotranslational misassembly.

Overall, this body of work strongly supports the concept that homomers frequently assemble cotranslationally, and that this has imposed constraints on protein evolution.

## Results

### Homomeric interface residues have a C-terminal bias

We hypothesized that a way of minimizing the potential for misassembly would be to localize homomeric interfaces towards the C-termini of polypeptide chains. Therefore, we examined the locations of interfaces in a large set of non-redundant homomer structures.

Figure 1A shows the relative enrichment of residues forming homomeric interfaces from N- to C-termini, considering all proteins in our dataset (see Supplementary Methods, “Structural analysis of interface location”). Strikingly, there is a highly significant greater tendency for interfaces to be formed by residues in the C-terminal halves of proteins. Overall, there is an 11.2% greater chance that a given point on a protein’s surface will be involved in a homomeric interface if it is located on the C-terminal half of the protein relative to the N-terminal half. This trend is also conserved across evolution: of the 16 out of 17 species with at least 100 non-redundant homomer structures in our dataset have positive interface enrichments in their C-terminal halves (Figure 1B). Furthermore, when bacterial or eukaryotic complexes are considered collectively, a significant enrichment in C-terminal interface residues is also conserved for each group (Figure S1).

Overall, this analysis is strongly supportive of the hypothesis that there has been evolutionary selection for homomer interfaces to be localized towards C-termini. However, the magnitude of this trend can be explained by evolutionary selection in only a proportion of individual homomeric polypeptide chains. For instance, amongst the 4834 bacterial homomers for which structures are available covering full-length polypeptide chains, 2722 (56.3%) have larger homomer interface area on their C-terminal half than their N-terminal half. While this is highly significant, it could also be explained by evolutionary selection in just 12.6% of homomers. It is possible that other mechanisms could act to minimize the harm of cotranslational misassembly, such as internal mechanisms like posttranslational folding or a linker separating the homomeric interface and the rest of the protein, or external mechanisms such as chaperones.

Importantly, this bias towards C-terminal interface residues was not observed for amino acids in the interfaces of heteromeric complexes, in archaea, bacteria, and eukaryotes (Figure S1C-E). This suggests that it relates to a specific evolutionary pressure relevant only to homomers. Our hypothesis is that the C-terminal bias of interface residues in homomers is a way of preventing cotranslational (mis)assembly driven by the high local concentration of unfolded nascent chains on the polysome.

### Cotranslationally aggregating homomers have larger N-terminal interfaces

We carried out an *in vivo* image-based high-throughput screen with a set of 611 native *E. coli* homomers. (Please refer to Figure S2 for a flow chart of the methodology). A GFP reporter domain was fused to the C-terminus of each homomer, and a supervised machine-learning approach was applied to automatically analyze the images of around one thousand cells for each protein. The fluorescence intensity and the intracellular localization of the fluorescent signal indicate the stability of the homomer. Overall, three distinct states were observed (Figure S2 and S3):

i. ells with homogeneously high GFP signal throughout the cell, indicating that the protein is folded and soluble, which we will refer to as ‘Green cells’.
ii. Cells with detectable fluorescent Inclusion Bodies (IB), which we will refer as ‘IB cells’. The presence of an IB detectable by flouresence microscopy is an indication that the protein is expressed and the GFP portion at the C-terminus is folded and fluorescent, so aggregation is likely to take place post-translationally (de Groot et al., 2008).
iii. Cell with GFP signal at background levels, which we will refer as ‘Dark cells’. These cells may indicate on one of two scenarios: either the protein has a low basal expression level, or the protein aggregates before it has fully folded. As GFP folds cotranslationally (Ugrinov and Clark, 2010), a lack of GFP signal means that aggregation is likely to take place cotranslationally. To rule out low basal expression, we compared our results with a recent study that used the same library to investigate the overall solubility of *E. coli* proteins during *in vitro* translation (Niwa et al., 2009). Comparing the two datasets allowed us to verify that all 611 homomers are indeed expressed *in vitro*, and the homomers with no GFP signal in our study are highly enriched in aggregating proteins *in vitro* (*p-value* 4.9×10^-5^ Figure S3). Therefore, it is likely that any absence of GFP signal in our *in vivo* screen also arises from cotranslational aggregation in living *E. coli* cells.

We allocated each homomer into one of the three groups (for details see Materials and Methods), and asked if high aggregation tendency correlates with N-terminal enrichment of homomeric interface residues, which may explain the bioinformatic analysis above (Figure 1 and S1). About a fifth of the homomers exhibited a preponderance of 'Dark cells', and these are significantly enriched in N-terminal interface residues as compared to the soluble homomers, *i.e.* “Green cells”, which have more interface residues in the C-terminal halves of the chains.

### The position of the oligomerization domain determines assembly and solubility

The computational analysis showing an enrichment in homomeric interfaces towards the C-terminal halves of proteins, and the results of the *in vivo* screen above are supportive of the occurrence of cotranslational (mis)assembly. To further probe cotranslational (mis)assembly and to achieve a spatiotemporal understanding of these events, we engineered two constructs that are identical in sequence, but differ in the order of their domains. Both constructs contain an oligomerization domain, *i.e.* p53Tet, that is connected to YFP by a short linker (SL). The final polypeptide product has the same quaternary structure (Figure S5). One construct, *i.e.* p53Tet-SL-YFP, has the Tet domain at the N-terminus of the protein, thus allowing, at least in theory, cotranslational assembly. The other construct, *i.e.* YFP-SL-p53Tet, does not, because the Tet domain is at the C-terminus and will be translated last.

Using confocal microscopy, we observed a significant difference in fluorescence levels between the two constructs (Figure 1E), which is remarkable considering that they are identical in their sequence composition and quaternary structure. We confirmed the total protein expression and protein solubility using Western blotting (Figure 1F). In agreement with the microscopy data, only YFP-SL-p53Tet showed a band in the soluble fraction, which indicates its folded state. The p53Tet-SL-YFP protein extract was found only in the insoluble fraction. Since the total protein expression levels of both samples were the same, we can conclude that the difference in fluorescence is not due to events prior translation.

We then compared the fluorescence intensity of the two constructs using flow cytometry. We found that with the tetramerization domain at the C-terminus, the fluorescence is over an order of magnitude higher, which is in the range of the difference in fluorescence observed by microscopy between the ‘Green’ and ‘Dark cells’ groups (Figure S3). Similarly, using a Tet mutant which is dimeric rather than tetrameric (Gaglia et al., 2013; Lomax et al., 1998), we see the same phenomenon (Figure S4), suggesting that the misassembly rates in these constructs are a function of assembly *per se* rather than a specific oligomeric state. These engineered constructs provide evidence that homomer solubility can be a function of the interface-residue position along the sequence.

Interestingly, co-expression of the 30aa p53Tet peptide reduces misassembly (Figure S6). The expression of p53Tet-SL-YFP, with or without the peptide, was normalized to a monomeric variant that does or does not co-express the peptide, respectively. The rapid association-dissociation kinetics (Rajagopalan et al., 2011) of the tetrameric variant and the high expression (Natan and Joerger, 2012) of the p53Tet peptide [not the case for the p53 full-length protein (Natan et al., 2011)]can explain this rescue, likely by masking of the homomeric interface of the p53Tet-SL-YFP polypeptide by the p53Tet peptide to prevent misassembly.

### Extending linker length reduces misassembly

We next sought to assess the effect of the addition of a long and flexible linker to the fate of the protein’s stability under the reasonable assumption that increasing the distance between the oligomerization domain and the reporter will reduce misassembly. We based our assumption on the fact that when fully-exposed from the ribosome tunnel, the flexibility of the linker (in comparison to that of the cotranslational folded reporter) will determine its relative concentration, *e.g.* Movie S1 *vs.* S3. More significant factor would be the consequence after assembly, meaning that the reporter’s nascent-chains are physically apart as a function of linker length, with a soluble domain, *i.e.* the linker, separating to determine the degree of misassembly (Figure 1A).

Three different linkers were used. Firstly, a short linker (SL), which is a five amino acid (aa), glycine-based linker. Second, a medium and long linker composed of one or two copies of the lipoyl domain of the dihydrolipoyl acetyltransferase enzyme (Jones et al., 2000). The medium-linker (ML) is 50aa long, comprised of one lipoyl domain from *B. stearothermophilus*, and the long-linker (LL) is 100aa long, representing this lipoyl domain plus another lipoyl domain from *E. coli* (Lengyel et al., 2008; Radford et al., 1989).

We also introduced a control construct, which is a monomeric variant of the Tet domain with a single point mutation (Figure 2A). Thus we can calculate the ratio of fluoresce intensity between the tetrameric *versus* monomeric variants to quantify the contribution of homomerization to misassembly.

**Figure 2.**
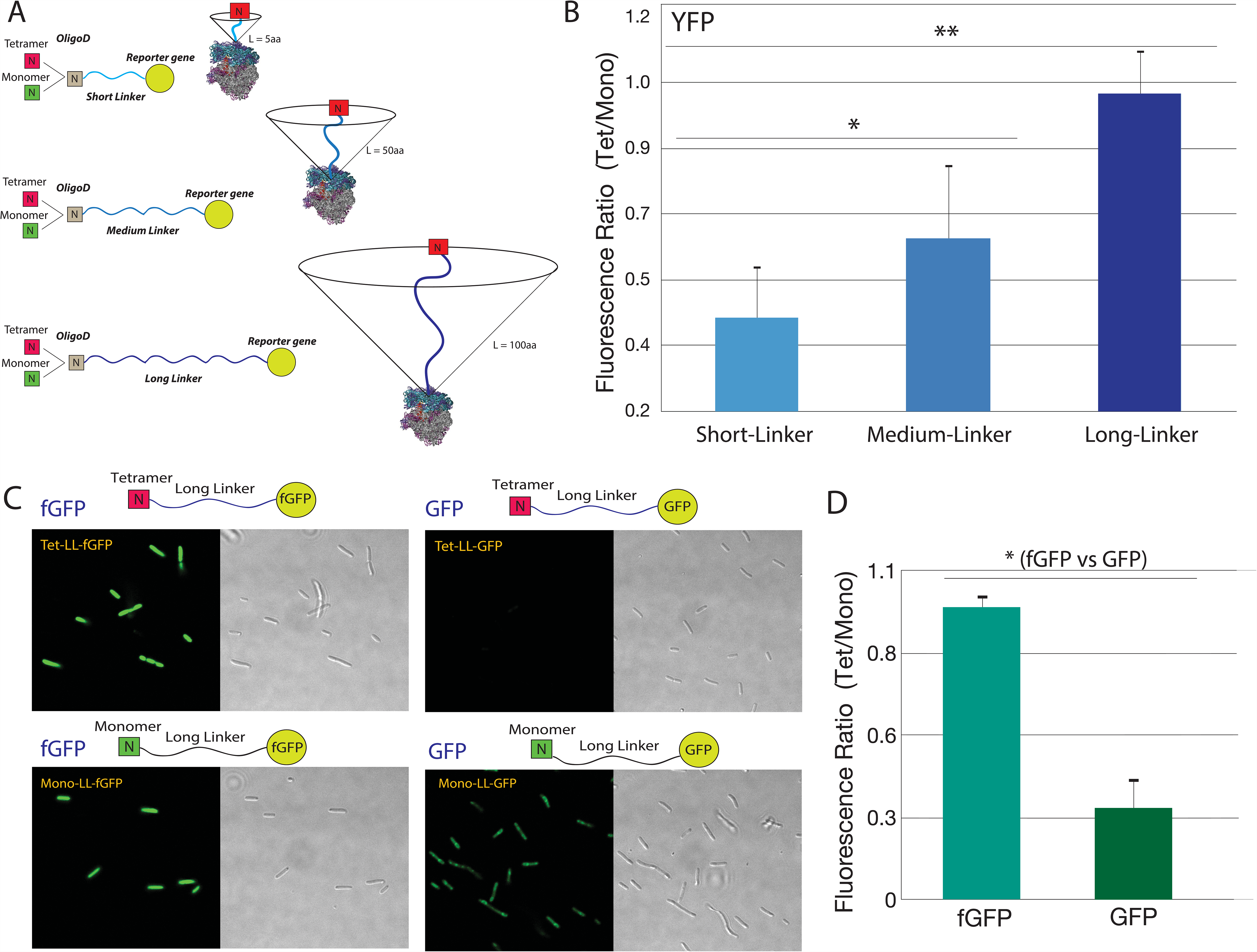
Extending the linker increases correct folding and assembly. (A) Scheme of the different constructs. All constructs have an oligomerization-domain at the N-terminus. The comparison is between tetrameric and monomeric variants, which differ by a single amino acid. The medium- (ML), and long (LL) linkers are one, and two flexible lipoyl domains, respectively. On the right is an unscaled diagram of the nascent-chain ribosome complexes after the translation of the linker and before the reporter exiting the ribosome tunnel. The cone schematically represents the volume p53Tet can diffuse throughout, and thus the relative distance between nascent chains and their local concentration, which are factors that may influence misassembly. (B) Flow Cytometry analysis provides us with the ratio of the fluorescence of tetrameric-to-monomeric variants, and this increases with linker length. This indicates fewer cotranslational misassembly events. This is either due to the dilution of the oligomerization domain in the confined environment of the polysome and hence a lower local concentration, or due to presence of the linker as a soluble barrier between the tetramerisation domain and the beta barrel domains of the reporter. (C) Confocal microscopy images of constructs with fGFP or GFP reporter genes. Similarly to the long-linker YFP variants, fGFP shows high fluorescence without a significant difference between the monomeric and tetrameric variants (no saturation was allowed). For the GFP variants, a strong effect is observed under the same conditions. (D) Flow Cytometry analysis of the ratio of tetrameric to monomeric variants for each reporter gene. The data presented is the average of these ratios as measured on different days. As seen by microscopy images, no significant difference was observed between the fGFP varinats. However a 3.5-fold higher fluorescence was observed for the GFP monomeric compared to tetrameric varinats [*p-value* *<0.05, ** <0.01 (see also Figure S4 for YFP)]. The beta-barrel structures were adapted from the Protein Data Bank, entry 3MGF.

Importantly, the three monomeric constructs showed a similar level of fluorescence (Figure S7). In contrast, the fluorescence intensity of the tetramer constructs increases with increasing linker length. The ratio of the fluorescence of the tetramer-to-monomer strains of each linker showed a positive correlation between the length of the linker and the extent of correct assembly of the protein (Figure 2B). These results suggest that the increase in the linker length leads to more efficient assembly. This could be due to a decrease in cotranslational interactions between the nascent chains on the polysome (through the in the radius of gyration of the nascent chains, and thus a decrease in local concentration) (Ruff et al., 2015). Alternatively if there is cotranslational assembly, the longer distance between the assembly-site and newly-exposed reporter will reduce the likelihood of nonspecific interactions and misassembly, and support more efficient co-translational assembly (Figure 1A).

### Fast folding of the reporter promotes efficient assembly

The balance between translation and folding rates is crucial for the fate of synthesized proteins. For example, changes in translation rates *via* a small number (Pechmann and Frydman, 2013) or even a single (Tsai et al., 2008) synonymous substitution of a rare to abundant tRNA codon changes the translation-folding balance and affects folding efficiency (O'Brien et al., 2012b). This is because slower translation rates provide a longer time for folding to occur. The fine balance between translation and folding rates has also been shown to determine the final tertiary structure of the beta-barrel cotranslationally folding fluorescent protein family which includes GFP (Sander et al., 2014).

To examine the role of protein folding rate in misassembly, we used two monomeric GFP variants with different folding rates: a fast folding GFP variant (fGFP) and a wild-type like folding variant (GFP) (Xu et al., 2013). In order to isolate the effect of folding-rate, we engineered constructs with long linkers. For constructs with a long linker and fast-folding reporter gene, *i.e.* YFP, we show that both the tetrameric and monomeric variants have similar fluorescence intensity (Figure S7). With this as a baseline, we then tested the GFP variant constructs. As before, all results were normalized for expression levels by quantifying the fluorescence ratio between strains that express tetrameric or monomeric variants. Using confocal microscopy, we observed that both monomeric and tetrameric fGFP variants presented similarly bright fluorescence levels (Figure 2C). This is similar to the results for YFP, which is not surprising as fGFP and YFP share the three fast folding mutations (F64L, V68L, S72A) located at the center of the beta-barrel (Sheff and Thorn, 2004; Xu et al., 2013).

In contrast, the slow folding GFP (GFP) showed a significant difference between the monomeric and tetrameric variants (Figure 2C). To quantify these observations, we used flow cytometry under the same experimental conditions. While the monomeric and tetrameric fGFP variants have essentially the same fluorescence levels, the tetrameric GFP has ~3.5-fold lower fluorescence than the monomeric GFP variant (Figure 2D). Interestingly, by cultivating the cells with the tetrameric and monomeric GFP variants at 18°C, the observed difference was reduced significantly (Figure S8).

Luciferase (Luc) is a two-domain, slow folding protein, with a completely different architecture to the beta barrel proteins. We cloned a Luciferase sub-library with both short and long linkers (Table S1). The *in vivo* results show similar trends to the other reporter genes (Figure S9A). The short linker monomeric construct had a much stronger signal than that of the tetrameric variant. While extending the linker length caused an increase in total fluorescence, the long linker monomeric variant still exhibited greater fluorescence than its tetrameric counterpart. This confirms that, analogous to the GFP constructs, a slow folding reporter domain provides ample opportunity for misassembly, even with a long linker.

### Misassembly and its recovery using an *in vitro* translation system

To further investigate whether this phenomenon is cotranslational, we expressed the constructs in a well-characterized expression system, *i.e.* the PURE *in vitro* translation system. This methos is robust, well-established that contains only the minimal components required for translation (Shimizu et al., 2001; Shimizu et al., 2005). This defined system allows us to examine biochemically whether our *in vivo* observations are due to a defect in cotranslational assembly. By controlling the [mRNA: Ribosomes] ratio, the equilibrium can be shifted from monosomic to polysomic translation (Figure 3B). By doing so, one can control the probability of two nascent chains interacting cotranslationally.

**Figure 3.**
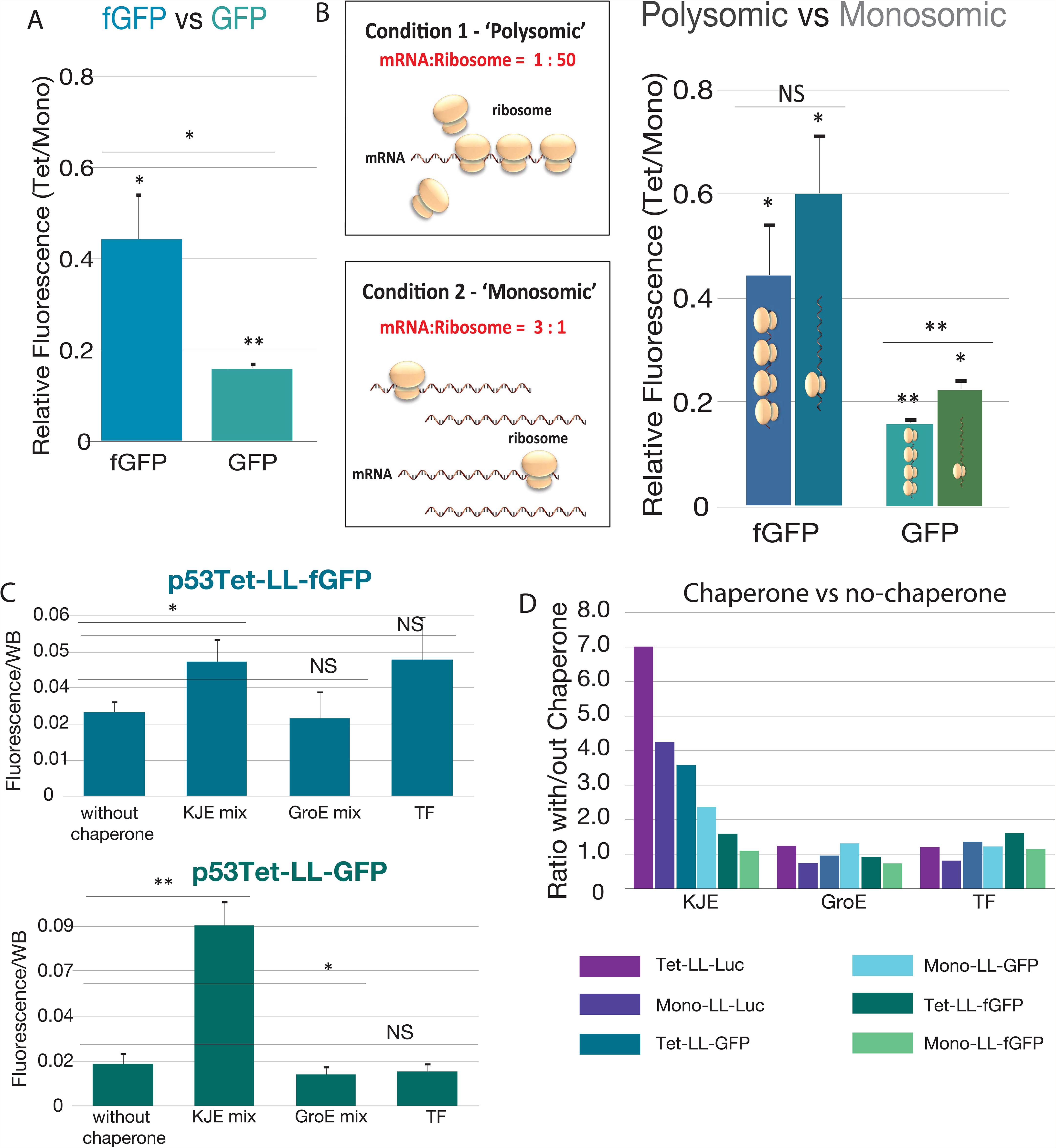
*In vitro* translation using the PURE system shows that misassembly takes place cotranslationally and is affected by the chaperones DnaKJ/GrpE. (A) Using the well-defined PURE translation system we measured the differences between the tetrameric and monomeric variants of fast folding (fGFP) and slow folding GFP. Correct folding of each construct was calculated as the ratio between the fluorescence signals and total protein as established by WB (for example see Figure S8). The value of the tetrameric construct was then divided by the monomeric constructs of the same reporter gene, *i.e.*, fGFP or GFP. (B) Similarly to the *in vivo* results, misassembly was dependent on the oligomerization-domain and folding rate. Moreover, by tuning the Ribosome:mRNA ratio we could influence the probability of the nascent chains interacting, *i.e.* polysomic conditions with 3:1 mRNA:Ribosome or monosomic conditions with 1:50 mRNA:Ribosome. The GFP tetrameric variants showed a decrease in solubility in comparison to the monomeric variant; a difference that is reduced if the conditions are shifted towards monosomes. fGFP variants show the same solubility in both polysomic and monosomic conditions. The same experiments were conducted using Luciferase (Luc) as the reporter gene. Luc is a larger protein than GFP and has a slower folding rate. For Luc, similar trends were observed as for the slow folding GFP. For Luc, there were greater differences between tetramer and monomer, between the polysomic *versus* monosomic conditions, and with and without chaperones. The results are summarized in Figure S7. (C) Slow and fast GFP folding, p53Tet-LL-GFP and p53Tet-LL-fGFP, respectively were tested using three chaperone groups. The first is the KJE mix, which includes DnaK, DnaJ and GrpE. The second is the GroE mix, which includes GroEL and GroES, and the third is Trigger Factor (TF). Interestingly, only slow folding GFP was affected by chaperones, and only by the KJE mix. (D) Summary of the effect of the different chaperones on GFP, fGFP and Luc sub-libraries, presented as the ratio of with/without chaperone, measurmenTs were repeated at least three times (see Suppl Methods). Overall, the effect of chaperones correlated with oligomeric state, *i.e.* tetramer *versus* monomer and with folding rate, *i.e.*, fast- and slow-folding proteins. The highest rescue effect was achieved by the KJE mix, particularly with the tetrameric slow folding Luc and GFP. All experiments are summarised in Figure S13 and in Suppl Methods. [*p-value* * < 0.05, ** < 0.01, NS = Not Significant].

In summary, our results using the PURE system at 37°C are in agreement with the *in vivo* measurements. We first examined the fGFP and GFP sub-libraries with a long linker. The levels of correctly folded reporter translated at a low [mRNA:Ribosomes] ratio of 1:50 were quantified by the ratio of fluorescence (or luminescence) to the expression level of the fully translated polypeptide, as measured by Western blot (Figure S10). The monomeric variant of fGFP had ~2-fold higher fluorescence than the tetrameric variant, while the monomeric variant of GFP was ~6-fold higher than its tetrameric counterpart (Figure 3A).

To further examine the significance of the folding rate of the reporter, we also analyzed the Luciferase sub-library (Figure S9B) and YFP sub-library (Figure S11). Luciferase presents similar trends, but with larger differences than the YFP or fGFP sub-libraries, in agreement with the *in vivo* results. Moreover, and also in agreement with the *in vivo* results, the short-linker Luciferase sub-library showed the lowest signal for the p53Tet-SL-Luc variant, and a significantly higher signal for the p53Mono-SL-Luc variant. Upon extension of the length of the linker in the Luciferase reporter, the difference between the monomeric and tetrameric variants was still significant.

Interestingly, extending the length of the linker has a different level of impact *in vitro versus in vivo*. For the sub-libraries that contains Luciferase and YFP reporters, the observed ‘rescue’ (as calculated by the ratio of Tetramer/ Monomer) is more significant *in vivo* than *in vitro*. By extending the linker, the tetrameric variant is more likely to assemble correctly (like the monomeric variant) *in vivo* but less so *in vitro*. The reason may be either the slower translation-rate reported *in vitro*, or the higher macromolecular crowding *in vivo* in comparison to the more dilute environment of the PURE system. The results may also reflect on the presence of specific proteins, *e.g.*, chaperons as also examined inhere. Nevertheless, these results underline the differences between *in vivo* and *in vitro* systems.

By decreasing the [mRNA:Ribosome] ratio by 150-fold (to 3:1) the probability of polysome formation is drastically reduced. This in turn reduces the probability for two nascent chains to assemble cotranslationally. When comparing the fGFP, GFP and Luciferase sub-libraries under monosomic and polysomic conditions, there is a clear trend supporting the hypothesis that a lower local concentration of nascent chains, as in the monosomic conditions, can rescue slow folding reporters from misassembly (Figure 3B and S9). The tetrameric fGFP constructs (p53Tet-LL-fGFP) showed only a marginal difference between the two conditions, and no difference was observed for the other fGFP constructs. As expected, and in contrast to fGFP, the sub-library of GFP showed a significant difference between the polysomic and monosomic conditions. The most significant recovery was observed for the tetrameric variant, as expected.

The Luciferase sub-library was also consistent with these observations, and similar to the GFP sub-library (Figure S9C). As with the other sub-libraries, the largest difference was between the polysomic and monosomic conditions that expressed p53Tet-LL-Luc and p53Tet-SL-Luc, with 5-fold higher misassembly in polysomic conditions.

In summary, the biochemically defined PURE system allowed us to tightly regulate the mRNA-to-ribosome ratio, and thus achieve polysomic and monosomic conditions *in vitro*. These observations are in agreement with the *in vivo* findings: the factors that favour efficient assembly are the C-terminal position of the Tet, long linkers and a fast folding reporter domain. For all these factors, misassembly is reduced in the monosomic condition, where there is a lower local concentration of nascent chains, compared to the polysomic condition (summarized in Figure 5).

### The effect of chaperones on misassembly

The selectivity of the PURE system allows us to test the effect of different chaperones on rescuing constructs from misassembly (Figures 3 and S7). We tested three chaperone groups. The first, “KJE mix”, includes DnaK, DnaJ and GrpE. The second is the “GroE mix”, which includes GroEL and GroES, and the third is Trigger Factor alone (TF).

Several conclusions could be drawn from the results:

i. The overall profiles of the effect of the chaperones are similar for the two structurally distinct slow-folding proteins, *i.e.*, GFP and Luc. Remarkably, the chaperone effect on GFP and Luc is much more alike than GFP and fGFP, which share >95% sequence similarity.
ii. As stated previously in the literature, *in vivo* aggregation assays show that homomers require DnaK for cotranslational folding (Calloni et al., 2012). Interestingly, the KJE chaperone mix (DnaK, DnaJ and GrpE) had the largest overall rescue effect, and it correlates with the proteins’ folding rate, as well as their oligomeric state.
iii. The GroEL mix only had an effect on the tetrameric variants with relatively slow folding, *i.e.* p53Tet-LL-GFP and p53Tet-LL-Luc, but not their corresponding monomeric variants. This is in agreement with previous work showing that reactivation of (monomeric) Luc was observed with a KJE mix, but not with GroEL and GroES (Jaenicke, 1991) (Niwa et al., 2012).
iv. TF did not have a drastic effect in our system, consistent with previous observations in the literature (Shieh et al., 2015). The ribosome-associated chaperone trigger-factor interacts directly with nascent polypeptide chains as they emerge from the ribosome exit tunnel (O'Brien et al., 2012a), and small protein domains can fold under the “cradle” created by TF, which may explain our results.

In summary, the chaperone DnaKJE seems to mitigate the effect of homomeric interactions on misassembly in this *in vitro* system. This provides at least a partial explanation for why homomeric contacts are tolerated in N-terminal positions in many naturally occurring proteins in organisms.

### *In silico* simulations visualize cotranslational assembly

To estimate the probability of nascent chain interactions occurring in the context of polysomes, and to gain insight into the mechanism of cotranslational assembly at atomic detail, we carried out *in silico* simulations of translation, folding and assembly. We used coarse-grained residue-level Brownian-dynamics simulations for three representative constructs with the YFP reporter. We focused on an inter-ribosomal geometry identified in tomographic reconstructions of experimentally determined *E. coli* ribosomes (Brandt et al., 2009) which has two peptide exit tunnels in close proximity. Using this model, we observed cotranslational folding and assembly, posttranslational assembly as well as simulations where no assembly occurs (Figure 4 and Movies S1-S3).

**Figure 4.**
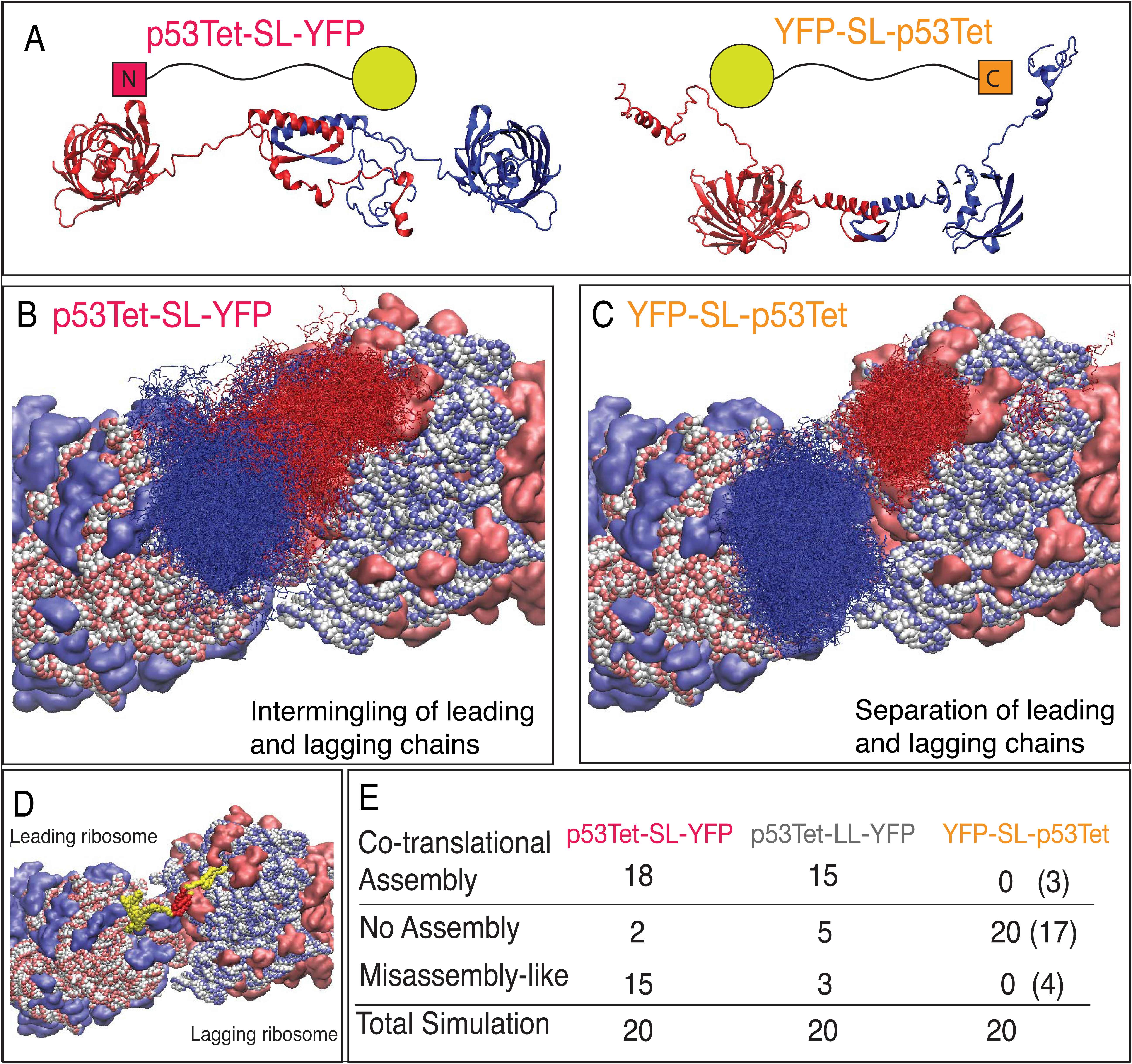
*In silico* simulation of translation of different constructs. (A) Simulation snapshot of cotranslational folding of two neighboring nascent chains of p53Tet-SL-YFP and YFP-SL-p53Tet. Composite plot showing regions typically sampled by the two nascent p53Tet-SL-YFP chains up to the point at which the translation of the first chain was completed. The leading ribosome is shown on the left. (B) Snapshots of cotranslational events as captured by simulations of polysomic translation. Ribosomes: blue and pink represent positively and negatively charged amino acids, respectively. In the case of p53Tet-SL-YFP the two chains intermingle, as the oligomerization domains assemble cotranslationally. (C) Same as (B) but showing a typical result for the YFP-SL-p53Tet construct; no intermingling of the two nascent chains occurs because the two oligomerization-domains remain unassembled. (D) Simulation snapshot showing the cotranslational assembly of two neighboring nascent chains. The leading ribosome is shown on the left. Tet is in red and YFP in yellow. (E) Table showing the number of cotranslationally, or posttranslationally (in brackets) assembly, misassembly-like event and total simulations. The relative positioning of the two ribosomes as found previously (Brandt et al., 2009).

We found that in simulations of the ribosomal synthesis of tetrameric N-terminal constructs (p53Tet-SL-YFP), cotranslational assembly of the constructs occurred in 90% of the simulations. As a result of this high-frequency cotranslational assembly, intermolecular interactions of the YFP domains occurred in 75% of the simulations (see also Figure S12). These intermolecular interactions likely represent misassembly events that inhibit the developmet of the fluorescence of the naturally monomeric YFP domain.

Extending the linker connecting the Tet and the YFP, decreases misassembly-like events. The reason was not due to a decrease in cotranslational assembly events, which are similar to the short linker construct. These results correlated well with the *in vivo* and *in vitro* results, again highlighting the ameliorating role of the long linker between the oligomerization-domain and reporter domains, as a diluter of the local concentration of the domains.

The simulations of tetrameric C-terminal constructs (YFP-SL-p53Tet) showed much less frequent assembly events, in agreement with our experimental results. When assembly did occur, it was a posttranslational event, or occurred as the newly synthesized chains were in the process of diffusing away from the ribosome exit tunnels. As expected from our hypothesis, intermolecular interactions of YFP were also rare events. These results indicate a clear relationship between the positioning of the Tet domain and the likelihood of misassembly events preventing the reporter domain’s fluorescence.

It is interesting to note that the tomographic reconstruction of *E. coli* polysomes previously reported (Brandt et al., 2009), which lies at the basis of our molecular simulations, provides a structural scenario in which cotranslational interactions of nascent chains can occur.

### Homomer misassembly reduces fitness and may constrain protein evolution

To assess the degree of homomer misassembly on the global bacterial fitness of *E. coli*, we measured the real-time growth-rates of fluorescent strains expressing the YFP and GFP sub-libraries. We found that the growth rates of strains with p53Tet at N–terminus (p53Tet@N) are lower in comparison to p53Mono@N or p53Tet@C (Figure 5 and Table S5). As expected, the largest differences were observed for the slow-folding GFP sub-libraries, and slightly less so for YFP. For the tetrameric *versus* the monomeric fGFP the differences were insignificant. The fluorescence levels and ratios were similar when measured by flow cytometry and microscopy. These *in vivo* results suggest that misassembly represents a burden to the cell that has a direct effect on growth rate and hence cellular fitness.

**Figure 5.**
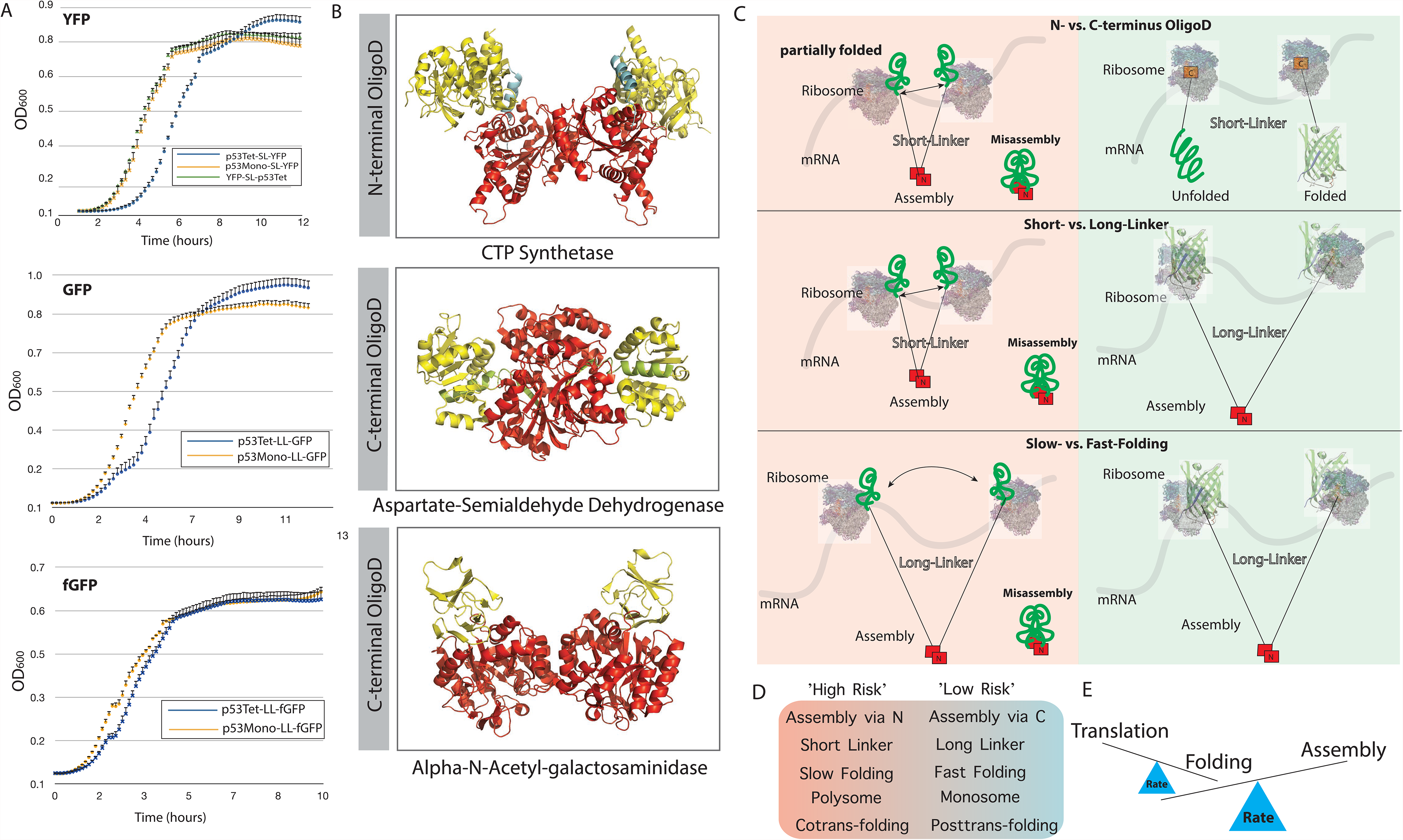
Cotranslational (mis)assembly and consequences. (A) Growth rate as a function of cotranslational assembly and misassembly. The sub-libraries of YFP, GFP or fGFP were expressed to measure growth in real-time. Each curve represents the average of same-day repeats. Measurements were repeated on different days, all showing the same trend, meaning N-terminal tetrameric constructs consistently grow more slowly than the other variants. (B) All *E. coli* homomeric protein structures were analyzed to create a library of protein structures with (i) a separate oligomerization-domain and (ii) with data to predict whether the protein folds co- or posttranslationally (O'Brien et al., 2012b). Three such protein structures were found. Oligomerization-domains are shown in red, domain-linkers in cyan, and other domains in yellow. CTP Synthetase PDB ID is 1s1m, and has an N-terminal oligomerization-domain, which may be compensated by the cyan linker. The other two proteins have C-terminal oligomerization-domains, no linkers, and are: aspartate-semialdehyde dehydrogenase (PDB ID 1t4b), and alpha-N-Acetyl-galactosaminidase (PDB ID 2p53). More information about these proteins is in Table S3. Summary (C-E): Once the amino acids that can generate a sufficient interface between the homomers fold, an assembly, whether steady or transient, may take place. According to the frequency and nature of that assembly, the proximity between the unfolded polypeptides will be determined and as such the probability of misassembly (left side). However, if an insufficient portion of the homomeric interface is generated, for instance if the oligomerization-domain is at the C-terminus (right side) or by increasing the linker length, misassembly is less likely to occur. An additional solution to misassembly would be to increase the rate of cotranslational folding. (D) Summary of possible mechanisms that enhance or prevent misassembly upon cotranslational folding. (E) This work can be viewed as an illustration of the balance between translation, folding and assembly. If sufficient folding occurs then assembly will follow. However if this assembly takes place while parts of the protein are still translating, *i.e.* unfolded, the balance will shift to misassembly.

Furthermore, in a protein complex immunoprecipitation (Co-IP) experiment in *E. coli* cultures we found a five-fold increase in the co-chaperone homologue Htp for the strain that expressed HA-tagged p53Tet-SL-YFP in comparison to YFP-SL-p53Tet (Table S4). This chaperone is the bacterial homologue of Hsp90, which buffers the effects of structurally destabilizing mutations (Hartl et al., 2011). This highlights the fact that misassembly imposes a burden on cells, and that the cell has evolved strategies to deal with this.

Given that misassembly can reduce cellular fitness, the C-terminal enrichment of homomeric interface residues we describe above may be the result of evolutionary selection against factors that promote misassembly (Figure 1). At the same time, there are multiple mechanisms that buffer against misassembly, some of which we investigated in this work. To explore this further, we studied a set of *E. coli* homomers that had both well-defined oligomerization domains as well as predicted co- or posttranslational folding signatures as calculated by the method in O’Brien *et al.* (O'Brien et al., 2012b). We were able to identify three such proteins (Figure 5 and Table S3) that represent homomeric full-length protein complexes in *E. coli*.

One of these three structures has an oligomerization-domain at the N-terminus, and in agreement with our prediction, the protein has a long linker right after the oligomerization-domain. Moreover, the oligomerization-domain is also predicted to fold posttranslationally (O'Brien et al., 2012b), which can provide an additional protection via temporal separation, *i.e.*, late assembly to avoid misassembly (Table S3). Neither of these factors holds for the other two *E. coli* homomers where the oligomerization-domain is at the C-terminus. With availability of new structures of large complexes, facilitated by developments in Cryo-EM, as well as functional genomics approaches to unravel the role of chaperones in cotranslational assembly, future studies will more accurately quantify the contribution of the different factors.

## Discussion

A protein’s amino acid sequence determines its structure, stability and interactions with other biomolecules. For homomeric proteins, which dominate protein quaternary structure space, the relationship between these parameters is only partially understood (Levy and Teichmann, 2013; Marsh and Teichmann, 2014). The stability of a monomeric protein, meaning its capacity to maintain the correct fold, is influenced by several factors, including the polypeptide’s capacity to fold cotranslationally, and its translation rate. Assembly, *i.e*. a protein’s native interactions with another identical chain to form a homomeric complex, will be driven, and determined by the availability and composition of its interface residues (Levy et al., 2012). However, as we hypothesized here, translation, folding and assembly of all the domains in the same polypeptide must be carefully balanced in space and time. The reason for this is that premature or incorrect assembly will impose a burden on the cell. To use an everyday analogy, in furniture production, there is a separation between manufacturing and assembly to avoid jumbling up the end product. Production and assembly of homomeric proteins follows the same logic. However, as proteins are synthesized in a highly crowded environment, the separation between the production and assembly needs to be enforced.

One way of achieving this is to position the residues mediating the assembly towards the end of the protein, so that it is synthesized before it starts assembling. Interestingly, it has been previously shown *in vitro* that refolding after denaturation of homomeric proteins is more challenging than for monomeric proteins due to premature assembly (Jaenicke and Lilie, 2000). This suggests that ribosomal protein synthesis may actually play a role in fine-tuning the correct assembly of homomers. In other words, these *in vitro* data may imply that for naturally occurring proteins, concurrent refolding and assembly has a higher misassembly rate than when folding occurs cotranslationally.

In order to investigate the conditions that promote misassembly, as well as its consequences, we established *in vivo*, *in vitro*, and *in silico* systems to examine diverse constructs of oligomerization and reporter domains. We found that misassembly can obstruct protein stability and decrease cellular fitness. At the same time, misassembly only occurs under certain conditions of oligomerization domain position, linker length, folding rate, *etc*. These factors are summarized in Figure 5.

The p53 oligomerization-domain (p53Tet) used here is known to fold cotranslationally (Nicholls et al., 2002). It is worth mentioning that cotranslational assembly leads to limited rotational freedom of the nascent chains in addition to an extreme increase in local concentration. Being linked to a second nascent chain at the N-terminus and the ribosome at the C-terminus, the protein’s capacity to ‘travel’ along the folding-landscape will be severely impaired.

Consistent results at the *in vivo, in vitro* and *in silico* levels allow us to put forward a spatiotemporal framework for correct translation, folding and assembly. First, the location of the oligomerization domain and used linker are both crucial: if the oligomerization-domain is N-terminal with a short linker, the reporter domains can be forced into close proximity directly upon translation on the polysome (Figure 1). Secondly, folding rate, which allows temporal separation of folding and assembly, may also play a role in safeguarding the stability of homomeric proteins.

These findings raise the question of what countermeasures and strategies exist in cells to cope with homomeric misassembly. While chaperones can serve as one solution, relying solely on chaperones would be a costly strategy. A more energy-efficient approach would be to avoid premature assembly in the first place, by evolving protein sequences that ensure a correct balance between translation, folding and assembly. The results shown here suggest that there has been significant evolutionary selection for the avoidance of homomer misassembly through the C-terminal localization of interfaces, but as we show using different chaperones, other mechanisms are likely to contribute.

Interactions between proteins are inter-molecular, stochastic events where the frequency and length of association are determined by the nature of the protein’s surface (Levy et al., 2012). In the confined environment of the polysome, assembly becomes an intra-molecular event that can compete with other intra-molecular events, *i.e.* folding. Moreover, a mutation that even weakly promotes a steady or transient interaction may have a significant effect on assembly. In other words, the cotranslational context may not only guide folding and assembly along the correct pathway as mentioned above, but also penalize any mutations that shift the equilibrium towards non-native interactions.

Our work contributes to the understanding of homomeric complexes, and the delicate balance between their translation, folding, and assembly as shaped by evolution. Moreover, this work suggests that for both native and non-native assembly sites, misassembly may take place, which leads to nascent chain aggregation and consequently to diseases. It is tempting to hypothesize that the disease associate genes of several neurodegenerative diseases are related to this type of misassembly. For example, the Huntingtin protein, which is believed to be the cause of Huntington’s disease (HD), has a N-terminal peptide with oligomerization capacity, and which accelerate significnalty amyloid formation (Tam et al., 2009). It may well be that cotranslational assembley as describe in this work, may accelerate amyloid formation in HD, as well as in other neurodegenerative diseases with assembely-like domain (Lashuel et al., 2013).

In summary, this work provides a new perspective on protein evolution and the cotranslational assembly of homomers, as well as a potential novel angle on the mechanism of protein aggregation in diseases.

## Contributions

The study was conceived and coordinated by EN and SAT.

The experiments were designed by EN, LHV, BK, BP, CP and PH.

The experiments were conducted by EN, TE, NS, AHE, BK, LD and PH. Bioinformatic analysis was conducted by TF and JAM, and simulations run by AHE. Machine learning analysis was conducted by PH.

Data analysis was conducted by EN, TE, AHE, TF, BK, GF, HP, BP CP and GC. The project was supervised by SAT, MMB, NS and CV.

The manuscript was drafted and refined by EN and SAT with contributions from all authors.

## Acknowledgment

We are grateful to C. Hyeon for the GFP unfolding illustration and A. Drummond for the generous gift of plasmids, and Günter Kramer and Bernd Bukau for their generous gift of Trigger Factor protein. We would also like to thank L. Byung-Gil for useful advice and to N. Sanchez De Groot for technical support. We thank C Vogel, MT Burgas and E Arbely for helpful suggestion and critical reading. MMB, TF and GC are supported by the Medical Research Council (MC_U105185859). TF is also supported by the Boehringer Ingelheim Fond. BP and CP would like to thank ‘Lendület’ Programme of the Hungarian Academy of Sciences and the Wellcome Trust for supporting this work, and the European Research Council (CP). PH would like to thank the National Brain Research Programme and the TEKES Finland Distinguished Professor Grant for their support. SAT thanks the Lister Institute, the MRC, the EMBL-European Bioinformatics Institute and the Wellcome Trust Sanger Institute. This work was supported in part by Grants-in-Aid for Scientific Research and MEXT (Japan)-Supported Program for the Strategic Research Foundation at Private Universities (2014-2019) and The Hirao Taro Foundation of KONAN GAKUEN for Academic Research. JM is supported by an MRC Career Development Award (MR/M02122X/1).

## References

Borgia, M.B., Borgia, A., Best, R.B., Steward, A., Nettels, D., Wunderlich, B., Schuler, B., and Clarke, J. (2011). Single-molecule fluorescence reveals sequence-specific misfolding in multidomain proteins. Nature 474, 662–665.

Brandt, F., Etchells, S.A., Ortiz, J.O., Elcock, A.H., Hartl, F.U., and Baumeister, W. (2009). The native 3D organization of bacterial polysomes. Cell 136, 261–271.

Calloni, G., Chen, T., Schermann, S.M., Chang, H.C., Genevaux, P., Agostini, F., Tartaglia, G.G., Hayer-Hartl, M., and Hartl, F.U. (2012). DnaK functions as a central hub in the E. coli chaperone network. Cell reports 1, 251–264.

de Groot, N.S., Espargaro, A., Morell, M., and Ventura, S. (2008). Studies on bacterial inclusion bodies. Future microbiology 3, 423–435.

Elcock, A.H. (2006). Molecular simulations of cotranslational protein folding: fragment stabilities, folding cooperativity, and trapping in the ribosome. PLoS computational biology 2, e98.

Gaglia, G., Guan, Y., Shah, J.V., and Lahav, G. (2013). Activation and control of p53 tetramerization in individual living cells. Proceedings of the National Academy of Sciences of the United States of America 110, 15497–15501.

Goodsell, D.S., and Olson, A.J. (2000). Structural symmetry and protein function. Annual review of biophysics and biomolecular structure 29, 105–153.

Hartl, F.U., Bracher, A., and Hayer-Hartl, M. (2011). Molecular chaperones in protein folding and proteostasis. Nature 475, 324–332.

Hartl, F.U., and Hayer-Hartl, M. (2009). Converging concepts of protein folding in vitro and in vivo. Nature structural & molecular biology 16, 574–581.

Iwasaki, S., and Ingolia, N.T. (2016). PROTEIN TRANSLATION. Seeing translation. Science 352, 1391–1392.

Jaenicke, R. (1991). Protein folding: local structures, domains, subunits, and assemblies. Biochemistry 30, 3147–3161.

Jaenicke, R., and Lilie, H. (2000). Folding and association of oligomeric and multimeric proteins. Advances in protein chemistry 53, 329–401.

Jones, D.D., Stott, K.M., Howard, M.J., and Perham, R.N. (2000). Restricted motion of the lipoyl-lysine swinging arm in the pyruvate dehydrogenase complex of Escherichia coli. Biochemistry 39, 8448–8459.

Kitagawa, M., Ara, T., Arifuzzaman, M., Ioka-Nakamichi, T., Inamoto, E., Toyonaga, H., and Mori, H. (2005). Complete set of ORF clones of Escherichia coli ASKA library (a complete set of E. coli K-12 ORF archive): unique resources for biological research. DNA research : an international journal for rapid publication of reports on genes and genomes 12, 291–299.

Lashuel, H.A., Overk, C.R., Oueslati, A., and Masliah, E. (2013). The many faces of alpha-synuclein: from structure and toxicity to therapeutic target. Nature reviews Neuroscience 14, 38–48.

Lengyel, J.S., Stott, K.M., Wu, X., Brooks, B.R., Balbo, A., Schuck, P., Perham, R.N., Subramaniam, S., and Milne, J.L. (2008). Extended polypeptide linkers establish the spatial architecture of a pyruvate dehydrogenase multienzyme complex. Structure 16, 93–103.

Levy, E.D., De, S., and Teichmann, S.A. (2012). Cellular crowding imposes global constraints on the chemistry and evolution of proteomes. Proceedings of the National Academy of Sciences of the United States of America 109, 20461–20466.

Levy, E.D., and Teichmann, S. (2013). Structural, evolutionary, and assembly principles of protein oligomerization. Progress in molecular biology and translational science 117, 25–51.

Lomax, M.E., Barnes, D.M., Hupp, T.R., Picksley, S.M., and Camplejohn, R.S. (1998). Characterization of p53 oligomerization domain mutations isolated from Li-Fraumeni and Li-Fraumeni like family members. Oncogene 17, 643–649.

Marsh, J.A., Rees, H.A., Ahnert, S.E., and Teichmann, S.A. (2015). Structural and evolutionary versatility in protein complexes with uneven stoichiometry. Nature communications 6, 6394.

Marsh, J.A., and Teichmann, S.A. (2014). Structure, Dynamics, Assembly, and Evolution of Protein Complexes. Annual review of biochemistry.

Mateu, M.G., Sanchez Del Pino, M.M., and Fersht, A.R. (1999). Mechanism of folding and assembly of a small tetrameric protein domain from tumor suppressor p53. Nature structural biology 6, 191–198.

Natan, E., Baloglu, C., Pagel, K., Freund, S.M., Morgner, N., Robinson, C.V., Fersht, A.R., and Joerger, A.C. (2011). Interaction of the p53 DNA-binding domain with its n-terminal extension modulates the stability of the p53 tetramer. Journal of molecular biology 409, 358–368.

Natan, E., and Joerger, A.C. (2012). Structure and kinetic stability of the p63 tetramerization domain. Journal of molecular biology 415, 503–513.

Nicholls, C.D., McLure, K.G., Shields, M.A., and Lee, P.W. (2002). Biogenesis of p53 involves cotranslational dimerization of monomers and posttranslational dimerization of dimers. Implications on the dominant negative effect. The Journal of biological chemistry 277, 12937–12945.

Nissley, D.A., and O'Brien, E.P. (2014). Timing is everything: unifying codon translation rates and nascent proteome behavior. Journal of the American Chemical Society 136, 17892–17898.

Niwa, T., Kanamori, T., Ueda, T., and Taguchi, H. (2012). Global analysis of chaperone effects using a reconstituted cell-free translation system. Proceedings of the National Academy of Sciences of the United States of America 109, 8937–8942.

Niwa, T., Ying, B.W., Saito, K., Jin, W., Takada, S., Ueda, T., and Taguchi, H. (2009). Bimodal protein solubility distribution revealed by an aggregation analysis of the entire ensemble of Escherichia coli proteins. Proceedings of the National Academy of Sciences of the United States of America 106, 4201–4206.

O'Brien, E.P., Christodoulou, J., Vendruscolo, M., and Dobson, C.M. (2012a). Trigger factor slows co-translational folding through kinetic trapping while sterically protecting the nascent chain from aberrant cytosolic interactions. Journal of the American Chemical Society 134, 10920–10932.

O'Brien, E.P., Vendruscolo, M., and Dobson, C.M. (2012b). Prediction of variable translation rate effects on cotranslational protein folding. Nature communications 3, 868.

Pechmann, S., and Frydman, J. (2013). Evolutionary conservation of codon optimality reveals hidden signatures of cotranslational folding. Nature structural & molecular biology 20, 237–243.

Radford, S.E., Laue, E.D., Perham, R.N., Martin, S.R., and Appella, E. (1989). Conformational flexibility and folding of synthetic peptides representing an interdomain segment of polypeptide chain in the pyruvate dehydrogenase multienzyme complex of Escherichia coli. The Journal of biological chemistry 264, 767–775.

Rajagopalan, S., Huang, F., and Fersht, A.R. (2011). Single-Molecule characterization of oligomerization kinetics and equilibria of the tumor suppressor p53. Nucleic acids research 39, 2294–2303.

Ruff, K.M., Harmon, T.S., and Pappu, R.V. (2015). CAMELOT: A machine learning approach for coarse-grained simulations of aggregation of block-copolymeric protein sequences. The Journal of chemical physics 143, 243123.

Sander, I.M., Chaney, J.L., and Clark, P.L. (2014). Expanding Anfinsen's principle: contributions of synonymous codon selection to rational protein design. Journal of the American Chemical Society 136, 858–861.

Sheff, M.A., and Thorn, K.S. (2004). Optimized cassettes for fluorescent protein tagging in Saccharomyces cerevisiae. Yeast 21, 661–670.

Shieh, Y.W., Minguez, P., Bork, P., Auburger, J.J., Guilbride, D.L., Kramer, G., and Bukau, B. (2015). Operon structure and cotranslational subunit association direct protein assembly in bacteria. Science 350, 678–680.

Shimizu, Y., Inoue, A., Tomari, Y., Suzuki, T., Yokogawa, T., Nishikawa, K., and Ueda, T. (2001). Cell-free translation reconstituted with purified components. Nature biotechnology 19, 751–755.

Shimizu, Y., Kanamori, T., and Ueda, T. (2005). Protein synthesis by pure translation systems. Methods (San Diego, Calif 36, 299–304.

Tam, S., Spiess, C., Auyeung, W., Joachimiak, L., Chen, B., Poirier, M.A., and Frydman, J. (2009). The chaperonin TRiC blocks a huntingtin sequence element that promotes the conformational switch to aggregation. Nature structural & molecular biology 16, 1279–1285.

Tsai, C.J., Sauna, Z.E., Kimchi-Sarfaty, C., Ambudkar, S.V., Gottesman, M.M., and Nussinov, R. (2008). Synonymous mutations and ribosome stalling can lead to altered folding pathways and distinct minima. Journal of molecular biology 383, 281–291.

Ugrinov, K.G., and Clark, P.L. (2010). Cotranslational folding increases GFP folding yield. Biophysical journal 98, 1312–1320.

Waldo, G.S., Standish, B.M., Berendzen, J., and Terwilliger, T.C. (1999). Rapid protein-folding assay using green fluorescent protein. Nature biotechnology 17, 691–695.

Wells, J.N., Bergendahl, L.T., and Marsh, J.A. (2015). Co-translational assembly of protein complexes. Biochemical Society transactions 43, 1221–1226.

Wright, C.F., Teichmann, S.A., Clarke, J., and Dobson, C.M. (2005). The importance of sequence diversity in the aggregation and evolution of proteins. Nature 438, 878–881.

Xu, C., Wang, S., Thibault, G., and Ng, D.T. (2013). Futile protein folding cycles in the ER are terminated by the unfolded protein O-mannosylation pathway. Science 340, 978–981.

